# Amino Acid Substitutions in the Na^+^/K^+^-ATPase May Contribute to Salinity Tolerance in Insects

**DOI:** 10.1101/2025.03.27.645771

**Authors:** Perla Achi, Victoria Iglesias, Preston Christensen, Rose C. Adrianza, Lanie Bavier, Robert Pena, Cullen McCarthy, Anil Baniya, W. Nate Collison, Simon C. Groen, Adler R. Dillman

**Author notes:** These authors contributed equally (Simon C. Groen, Adler R. Dillman). Lead contacts.

## Abstract

1. Environmental salinity levels vary naturally across terrestrial ecosystems but can be heightened locally by coastal proximity and desertification as well as human activities such as road salt application and agriculture. Since salt is essential for many physiological processes in insects, rising environmental sodium concentrations may drive behavioral changes, where insects select environments and food sources with suitable sodium levels, or evolutionary changes in constitutive or plastic physiological mechanisms to process salt, potentially altering ecological dynamics and species interactions.
2. Numerous hematophagous (blood feeding) insects such as the yellow-fever mosquito *Aedes aeqypti* are known to be able to breed in relatively saline environments. Among phytophagous (plant feeding) insects, grasshoppers can be important herbivores in arid and coastal salt-affected regions, whereas the monarch butterfly (*Danaus plexippus*) appears to perform relatively well on milkweed host plants growing in roadsides influenced by salt runoff. Several of these insects share a common trait: amino acid substitutions in the first extracellular loop of the Na+/K+-ATPase (NKA), a sodium pump crucial for maintaining ion balance. For the monarch these substitutions confer resistance to toxic cardenolides from milkweeds, but it is unclear whether NKA substitutions may influence salt tolerance.
3. Here, we investigate whether the NKA substitutions found in these insects may contribute to salt tolerance using gene-edited *Drosophila melanogaster* mutant strains as models. We show that flies with substitution Q111L (found in *Aedes* mosquitoes) or a combination of Q111L and A119S (found in grasshoppers) exhibited greater salt tolerance, whereas flies carrying the combination of substitutions found in the monarch (Q111V, A119S, and N122H) did not.
4. Our results suggest that the monarch may rely on alternate mechanisms for salt tolerance and that its NKA substitutions are important primarily for cardenolide resistance. However, substitution Q111L and the combination of Q111L and A119S may be relevant for salt tolerance in a variety of insects. Uncovering mechanisms of salt tolerance enhances our understanding of species distributions, ecological interactions, and evolutionary physiology in response to changing environmental salinity levels.

## Introduction

Salinity levels vary geographically across terrestrial ecosystems, particularly in coastal regions (Lorrain-Soligon et al., 2023). Sodium levels in soil range from < 0.05 % (8.5 mM) to > 1.75% (300 mM) in strongly saline areas (Department of Primary Industries and Regional Development, Western Australia, 2024). Many insect species—including phytophagous, hematophagous, and aquatic insects—may thus encounter a wide range of salt concentrations across their habitats (Santiago-Rosario et al., 2025). Sodium (Na^+^), while generally non-essential for most plants, is a key nutrient for animals to support development and physiological processes. Insects have therefore evolved several behavioral and physiological adaptations to not only ensure adequate sodium intake but also to process excess salt, as it may become toxic at levels higher than a physiological level of c. 0.9% (154 mM; Snell-Rood et al., 2014; Pontes et al., 2022; Santiago-Rosario et al., 2022). As an example of behavioral adaptation, butterflies engage in puddling, where they extract sodium from soil to compensate for the relatively low salt contents of their host plants, which is often < 0.05% (Xiao et al., 2010; Snell-Rood et al., 2014). This behavior is essential for supporting nervous system functions, growth, and reproductive processes (Lamie et al., 2025). Interestingly, plants may use sodium withholding as an anti-herbivore defense mechanism since they do not require high sodium for metabolic functions (Kaspari, 2020). On the other hand, following the “No-Escape-from-Sodium” hypothesis, sodium concentrations of plant tissues tend to reflect environmental salt levels (Santiago-Rosario et al., 2022). Sodium concentrations of salt-tolerant plants such as *Atriplex* saltbushes and salt-accumulating species such as the salt marsh grass *Spartina alterniflora* can reach well above physiological levels for animals (Santiago-Rosario et al., 2022; Vasquez et al., 2006). As such, environmental salt levels influence the phytochemical landscape, which in turn may affect interactions with phytophagous animals and even the structure of ecological communities (Santiago-Rosario et al., 2022).

Insect adaptations to fluctuating salt levels could become more important now that salinity levels are increasing to above physiological levels for animals across large areas due to climate change-associated desertification and rising sea levels as well as human activities such as road salt application and agriculture (Hassani et al., 2020; Kengne et al., 2019; Snell-Rood et al., 2014).

Understanding how some species can tolerate and have adapted to increases in sodium levels is crucial for explaining species’ ecological distributions and interactions.

Hematophagous insects such as mosquitoes, sand flies, and stable flies form an ecologically important guild and may encounter relatively high levels of salt through blood-feeding and when breeding in saline environments (Bradley, 1987; Yee et al., 2021). For example, some representatives of the mosquito genera *Aedes, Anopheles*, and *Culex* such as *An. merus* can thrive in saline habitats (Kengne et al., 2019; Yee et al., 2021), while *Ae. aegypti* and *Ae. albopictus* are expanding to coastal regions and are capable of breeding in environments with moderate to high salinity levels of up to 1.5% (Ramasamy et al., 2021).

Phytophagous insects might experience high salinity if they must navigate salt-affected soils in between bouts of feeding on plants or when attacking plants growing in saline environments along the coast or at roadsides. Monarch butterflies (*Danaus plexippus*) appear to potentially benefit from feeding as larvae on milkweeds (*Asclepias* spp.) in roadside environments (with leaf sodium concentrations closer to physiological levels for animals), since males show increased muscle mass and females exhibit greater neural investment (Snell-Rood et al., 2014; Hund et al., 2024). Both male and female monarchs accumulate relatively high sodium levels in saline environments, whereas this is only observed for males with other butterfly species (Santiago-Rosario et al., 2025). Grasshoppers, including the eastern lubber grasshopper *Romalea microptera*, can be important herbivores of plants such as *Spartina alterniflora*, which grows in coastal salt marshes and other salt-affected regions (Haines and Montague, 1979; Li et al., 2023; Vincent, 2006). Leaves of this plant can reach sodium levels of up to 3.5% (Vasquez et al., 2006). Physiological studies have shown that grasshoppers may tolerate levels of salinity reaching 11% (Dadd, 1961). Such observations raise the question of how these insects adapted to elevated environmental sodium levels.

The Na+/K+-ATPase (NKA) is a highly conserved sodium pump across animal species (Clausen and Poulsen, 2013). In insects, it plays a crucial role in regulating ion gradients, cell signaling, and stress responses (Groen and Whiteman, 2022). Several phytophagous and hematophagous insects that can be found in salt-affected environments, including the monarch butterfly, grasshoppers, *Aedes* mosquitoes, *Glossina* tsetse flies, *Phlebotomus* sandflies, and *Stomoxys* stable flies, evolved amino acid substitutions in the first extracellular loop of the NKA alpha subunit (Table 1; Dobler et al., 2012; Yang et al., 2019). Do these substitutions contribute to salt tolerance, given the sodium pump’s function in ion regulation? For monarchs, substitutions in the NKA facilitated specialization on milkweeds, conferring resistance to toxic cardenolides through target-site insensitivity. However, it remains unclear whether these substitutions have pleiotropic effects on salt tolerance. Establishing whether NKA substitutions contribute to salt tolerance in insects that frequently encounter saline environments may provide mechanistic insights into these insects’ distributions, behaviors, and ecological interactions.

**Table 1.** Amino acid sequences of the Na+/K+-ATPases of insects in the orders Orthoptera, Diptera, and Lepidoptera that have been found to show variation in their levels of salt tolerance. Sequences of the Na+/K+-ATPases of our study model Drosophila (*D. melanogaster*) are also given for the wild type (QAN) and gene-edited knock-in mutant strains (QSN, LAN, LSN, and VSH). The sequences at positions 111–122 of the Na+/K+-ATPase alpha subunit are shown (numbering relative to the pig ATP1A1 sequence).

| Order/Family | Species | Common Name | 111 | 112 | 113 | 114 | 115 | 116 | 117 | 118 | 119 | 120 | 121 | 122 |
| --- | --- | --- | --- | --- | --- | --- | --- | --- | --- | --- | --- | --- | --- | --- |
| **Orthoptera** |  |  |  |  |  |  |  |  |  |  |  |  |  |  |
| **Tetrigidae** | *Tetrix japonica* | Eastern common groundhopper | Q | A | S | T | V | E | E | P | A | D | D | N |
| **Romaleidae** | *Romalea microptera* | Eastern lubber grasshopper | L | A | S | T | V | E | E | P | S | D | D | N |
| **Acrididae** | *Locusta migratoria* | Migratory locust | L | A | A | T | V | E | E | P | S | D | D | N |
|  | *Schistocerca americana* | American bird grasshopper | L | A | S | T | V | E | E | P | S | D | D | N |
| **Diptera** |  |  |  |  |  |  |  |  |  |  |  |  |  |  |
| **Drosophilidae** | *Drosophila melanogaster* | Vinegar fly | Q | A | S | T | S | E | E | P | A | D | D | N |
| **Glossinidae** | *Glossina morsitans* | Tsetse fly | Q | A | S | T | S | E | E | P | S | D | D | N |
| **Muscidae** | *Stomoxys calcitrans* | Stable fly | Q | A | S | T | S | E | E | P | S | D | D | N |
| **Culicidae** | *Culex pipiens* | Common house mosquito | Q | A | S | T | V | E | E | P | A | D | D | N |
|  | *Culex quinquefasciatus* | Southern house mosquito | Q | A | S | T | V | E | E | P | A | D | D | N |
|  | *Aedes aegypti* | Yellow fever mosquito | L | A | S | T | V | E | E | P | A | D | D | N |
|  | *Aedes albopictus* | Asian tiger mosquito | L | A | S | T | V | E | E | P | A | D | D | N |
|  | *Anopheles coluzzii* | Malaria mosquito | Q | A | S | T | V | E | E | P | A | D | D | N |
|  | *Anopheles gambiae* | Malaria mosquito | Q | A | S | T | V | E | E | P | A | D | D | N |
| **Psychodidae** | *Phlebotomus papatasi* | Sand fly | Q | A | S | T | S | E | E | P | S | D | D | N |
| **Lepidoptera** |  |  |  |  |  |  |  |  |  |  |  |  |  |  |
| **Pieridae** | *Pieris rapae* | Small cabbage white | Q | A | S | T | V | E | E | P | A | D | D | N |
|  | *Danaus plexippus* | Monarch butterfly | V | A | S | T | V | E | E | P | S | D | D | H |
| **Study model** |  |  |  |  |  |  |  |  |  |  |  |  |  |  |
| **Drosophilidae** | *Drosophila melanogaster* | QAN | Q | A | S | T | S | E | E | P | A | D | D | N |
|  |  | LAN | L | A | S | T | S | E | E | P | A | D | D | N |
|  |  | QSN | Q | A | S | T | S | E | E | P | S | D | D | N |
|  |  | LSN | L | A | S | T | S | E | E | P | S | D | D | N |
|  |  | VSH | V | A | S | T | S | E | E | P | S | D | D | H |

We used *D. melanogaster* (Drosophila) to study gene-edited knock-in mutant strains with amino acid substitutions at sites 111, 119, and/or 122 of the NKA, reflecting substitutions found in hematophagous and phytophagous species (Groen and Whiteman, 2016; Karageorgi et al., 2019). Monarchs evolved substitutions Q111V, A119S, and N122H (VSH); *Aedes* mosquitoes evolved substitution Q111L (LAN); several hematophagous flies evolved substitution A119S (QSN); and grasshoppers evolved both Q111L and A119S (LSN; Table 1; Dobler et al., 2012; Yang et al., 2019). Salt tolerance of Drosophila mutants was compared to a control strain with the wild-type sequence (QAN). We aimed to determine whether these NKA substitutions could influence salt tolerance in insects.

## Methods

### Drosophila maintenance and culturing

Drosophila strains [QAN], [LAN], [QSN], [LSN], and [VSH] with or without substitutions at positions 111, 119, and/or 122 of the NKA alpha subunit in w[1118] background, were obtained from the Bloomington Drosophila Stock Center (Karageorgi et al., 2019). Flies were maintained in vials with food media (Genesee Nutri-Fly) in an incubator at 26 °C with a 12-hour light/dark phase.

### Salt tolerance assay with Drosophila

Ten males and ten females were put into individual vials containing 4 mL Genesee Nutri-Fly food for 3 days, allowing for reproduction. Ten first-instar larvae were then placed into vials containing different diets; 4 mL of regular Genesee Nutri-Fly media or media containing an additional 2% or 3% NaCl, which was mixed in before heating during food preparation. Wild-type larvae did not survive on food containing ≥5% NaCl. Three to five vials, containing ten first-instar larvae each, were used for each strain-by-diet combination per trial, depending on availability of larvae. Vials were placed randomly on racks with neighboring vials placed on the outer edges in an incubator with a 12-hour light/dark phase at 25 °C. Data was recorded daily over a 12-day period measuring the proportions of larvae that pupated and reached adulthood.

### Graphing and statistical analysis

Graphs were generated as survival curves in PRISM, displaying the mean and standard error of the mean (SEM). For statistical analyses a binomial generalized linear model (GLM) was fitted for each developmental stage transition, modeling survival (i.e., the number of individuals that survived out of the total tested) as a function of strain, treatment, and their interaction using the base R statistics package (version 4.3.1). The R package emmeans (version 1.11.1) was used to calculate estimated marginal means (EMMs) and perform post-hoc pairwise contrasts both between strains within each treatment and between treatments within each strain. We further used the log-rank (LR) Mantel-Cox test to perform pairwise contrasts and validate results from the GLM. Interpreting the results of these analyses conservatively, results were deemed significant when both analyses indicated p values <0.05. Figures were assembled using Adobe Photoshop.

## Results

### Substitutions in the NKA influence salt tolerance in Drosophila

First-instar Drosophila larvae were fed diets with varying salt concentrations and monitored for the proportion that pupated and eclosed (reached adulthood) successfully. We aimed to determine whether NKA amino acid substitutions found across insects could contribute to salt tolerance (Table 1). All fly strains were significantly affected by the 2% salt treatment, with fewer individuals pupating and reaching adulthood than at 0% salt (p<0.05). The 3% salt treatment further exacerbated these patterns for each strain (p<0.05).

However, in pairwise comparisons with the control fly strain QAN, several knock-in strains were significantly less affected by saline environments (Fig. 1). Strain LAN showed no significant difference in pupation compared to QAN at 0% or 2% salt but had a higher rate of pupation in a saline environment of 3% (p≤0.022). LAN began to show a significantly higher rate of eclosion than strain QAN at 2% salt and continued to show this pattern at 3% (p≤0.012 and p≤0.048, respectively). While strain QSN showed no significant differences in pupation or eclosion relative to QAN at any of the salt concentrations tested (Fig. 1), LSN had a significantly higher rate of eclosion than QAN at 2% salt (p≤0.012). VSH flies showed significantly fewer individuals pupating and reaching adulthood at 0% (p≤0.042), but no other significant difference was seen for VSH (Fig. 1). Taken together, our analyses found LAN to be the most salt tolerant strain, with strain LSN also having a better chance of reaching adulthood at a sodium level of 2% (Fig. 1).

**Fig. 1.**
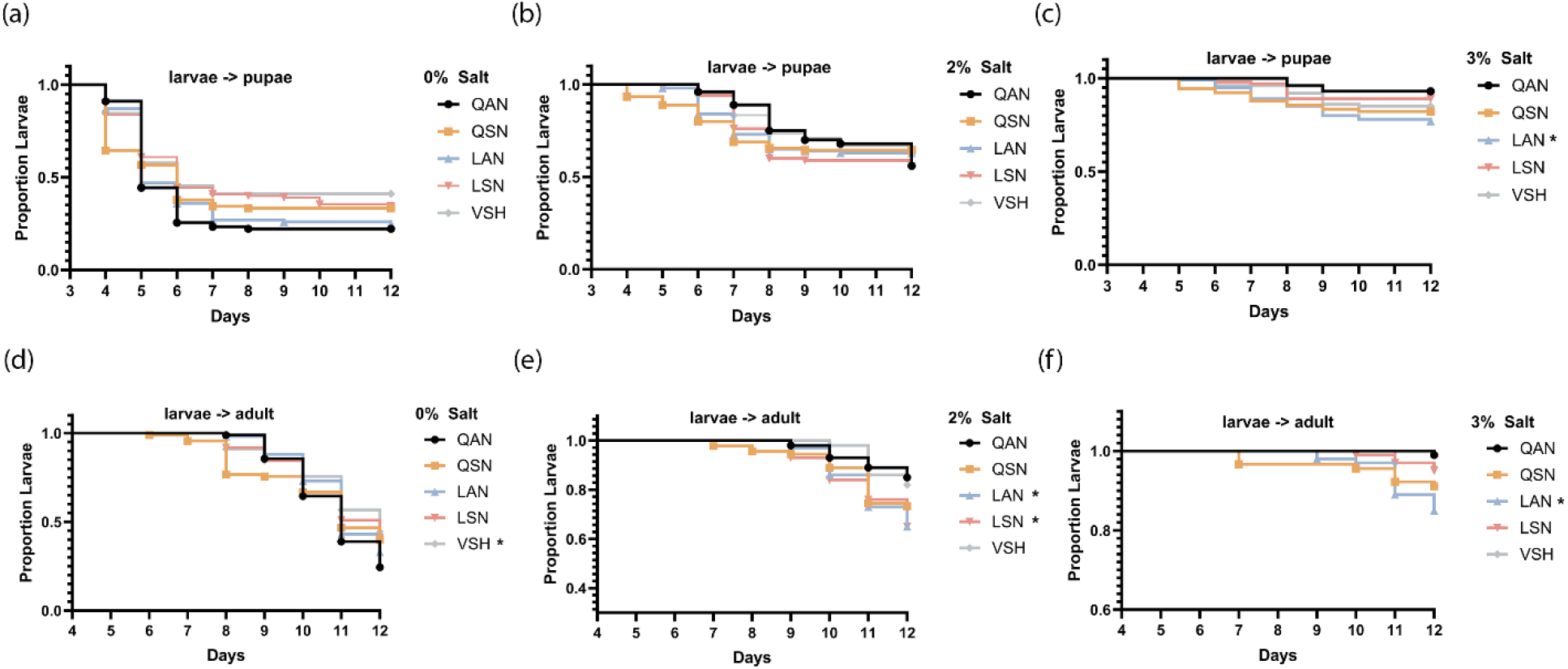
The LAN and LSN knock-in mutant Drosophila strains demonstrate higher salt tolerance by their increased ability of larvae to reach the pupal and adult stages. **A-C**. Proportion of first-instar Drosophila larvae of the QAN (wild type), QSN, LAN, LSN, and VSH strains that developed into pupae after being reared on diets containing 0%, 2% or 3% salt, respectively, over a 12-day period. **D-F**. Proportion of first-instar Drosophila larvae of the QAN (wild type), QSN, LAN, LSN, and VSH strains that successfully reached the adult stage post pupation after being reared on diets containing 0%, 2% or 3% salt, respectively, over a 12-day period. Data are represented as means ± SEM. Statistical analyses were performed using log-rank analysis and an analysis using a binomial generalized linear model with pairwise comparisons to the QAN strain.

## Discussion

Using gene-edited knock-in mutant Drosophila strains as models, we tested the effects of amino acid substitutions at sites 111, 119, and/or 122 in the first extracellular loop of the NKA’s alpha subunit—substitutions that evolved in a variety of phytophagous and hematophagous insect species known to occur in salt-affected environments. We hypothesized that substitutions in the sodium pump at these sites might influence salinity tolerance in insects.

Substitution Q111L (as observed in *Aedes* mosquitoes) or a combination of Q111L and A119S (as observed in grasshoppers) enhanced salt tolerance in Drosophila, with larvae of strains LAN and LSN showing higher pupation and/or eclosion rates on diets rich in salt. Previous research showed that wild-type (QAN) flies had significantly fewer larvae reaching adulthood at 1.75% salt and no survival at 5% salt (Riedl et al., 2016). We recapitulated these findings with diets containing 2% and 3% salt and observed that LAN flies tolerated these sodium levels significantly better than QAN in terms of reaching adulthood. In addition, strain LSN exhibited similar patterns of higher relative salt tolerance at 2% NaCl. This suggests that substitution Q111L and the combination of Q111L and A119S might improve insect salt tolerance.

Hematophagous insects may encounter salt in their environment in several ways, including by feeding on blood, which contains 0.9% salt in humans, and through aquatic breeding in arid or coastal salt-affected environments (Yee et al., 2021). The fact that *Aedes* mosquitoes evolved substitution Q111L may to some extent explain *Ae. aegypti*’s and *Ae. albopictus*’ successful range expansions into coastal areas (Ramasamy et al., 2021). However, we note that *Anopheles* and *Culex* also include species that occur in saline habitats but did not evolve Q111L (Dobler et al. 2012; Kengne et al., 2019), suggesting they evolved different salt tolerance strategies. Other hematophagous insects, including *Glossina* tsetse flies, *Phlebotomus* sandflies, and *Stomoxys* stable flies, can similarly be found in arid or coastal saline environments and evolved substitution A119S. However, since A119S did not confer enhanced salt tolerance, it is likely that these species evolved alternative mechanisms as well.

Phytophagous insects may primarily be exposed to elevated sodium levels through feeding on plants in salt-affected environments, such as roadsides and salt marshes (Snell-Rood et al., 2014). Several grasshopper species are successful herbivores of plants in coastal salt marshes and other salt-affected regions (Haines and Montague, 1979; Li et al., 2023; Vincent, 2006), can tolerate relatively high levels of salinity (Dadd, 1961), and evolved both substitutions Q111L and A119S. Our results for the Drosophila double mutant with these substitutions suggest that they may contribute to salt tolerance in these herbivores. Mechanisms of salt tolerance might become more important for insects since freshwater ecosystems are increasingly affected by salt contamination due to factors such as climate change-related sea-level rise and desertification as well as human activities, including groundwater extraction (Silver and Donini, 2021).

We hypothesized that insects with NKA substitutions linked to cardenolide resistance such as the monarch butterfly may experience pleiotropic effects. Leaves of milkweeds growing near roadways can contain elevated levels of Na^+^ and sodium accumulates to relatively high titers in both male and female monarch butterflies after having fed on such plants as larvae (Snell-Rood et al., 2014; Santiago-Rosario et al., 2025). However, our results indicate that the combination of substitutions Q111V, A119S, and N122H did not confer increased salt tolerance. This suggests that milkweed-produced cardenolides were likely to be more important agents of selection driving these NKA substitutions than environmental sodium (Karageorgi et al., 2019).

In summary, we found that substitution Q111L and the combination of Q111L and A119S may contribute to salt tolerance in insects. Their evolution could have enhanced physiological mechanisms for maintaining ion balance in a variety of phytophagous and hematophagous insect species. These contributions to salt tolerance may offer valuable insights into how insect species adapt to saline environments, influencing their ecological distributions.

## Acknowledgements

This study was supported by the National Institute of General Medical Sciences of the National Institutes of Health (award no. R35GM151194 to SCG and award no. R35GM137934 to ARD), the United States Department of Agriculture’s National Institute of Food and Agriculture (award no. 1032189 to ARD), and startup funds from the University of California Riverside (to SCG).

## Data Availability

The data that support the findings of this study are openly available at Mendeley Data: https://data.mendeley.com/datasets/2n2ttnpctw/1

## References

Bradley, T.J. (1987) Physiology of osmoregulation in mosquitoes. Annual Review of Entomology, 32, 439–462.

Clausen, M.J.V. & Poulsen, H. (2013) Sodium/potassium homeostasis in the cell. In: Metallomics and the Cell, 41–67, Springer.

Dadd, R.H. (1961) The nutritional requirements of locusts—V. Journal of Insect Physiology, 6, 126–145.

Department of Primary Industries and Regional Development, Western Australia (2024) Measuring soil salinity. Department of Primary Industries and Regional Development, Perth. Factsheet DPIRD-181. https://library.dpird.wa.gov.au/nrm_factsheets/26.

Dobler, S., Dalla, S., Wagschal, V. & Agrawal, A.A. (2012) Community-wide convergent evolution in insect adaptation to toxic cardenolides by substitutions in the Na,K-ATPase. Proceedings of the National Academy of Sciences of the USA, 109, 13040–13045.

Groen, S.C. & Whiteman, N.K. (2016) Using Drosophila to study the evolution of herbivory and diet specialization. Current Opinion in Insect Science, 14, 66–72.

Groen, S.C. & Whiteman N.K.. 2022. Ecology and evolution of secondary compound detoxification systems in caterpillars. In: Caterpillars in the Middle: Tritrophic Interactions in a Changing World, 115–163, Springer.

Haines, E.B. & Montague, C.L.. (1979) Food sources of estuarine invertebrates analyzed using ^13C/12^C ratios. Ecology, 60, 48–56.

Hassani, A., Azapagic, A. & Shokri, N. (2020) Predicting long-term dynamics of soil salinity and sodicity on a global scale. Proceedings of the National Academy of Sciences of the USA, 117, 33017–33027.

Hund, A.K., Mitchell, T.S., Ramιrez, M.I., Zambre, A., Hagg, L., Stene, A., Porter, K., Carper, A., Agnew, L., Shephard, A.M., Kobiela, M.E., Oberhauser, K.S., Taylor, O.R. & SnellRood, E.C. (2024) The potential of roadside verges as insect habitat: Road salt has few effects on monarch butterfly performance and migration. Conservation Science and Practice, 6, e13229.

Karageorgi, M., Groen, S.C., Sumbul, F., Pelaez, J.N., Verster, K.I., Aguilar, J.M., Hastings, A.P., Bernstein, S.L., Matsunaga, T., Astourian, M., Guerra, G., Rico, F., Dobler, S., Agrawal, A.A. & Whiteman, N. K. (2019). Genome editing retraces the evolution of toxin resistance in the monarch butterfly. Nature, 574, 409–412.

Kaspari, M. (2020) The seventh macronutrient: How sodium shortfall ramifies through populations, food webs and ecosystems. Ecology Letters, 23, 1153–1168.

Kengne, P., Charmantier, G., Blondeau-Bidet, E., Costantini, C. & Ayala, D. (2019) Tolerance of disease-vector mosquitoes to brackish water and their osmoregulatory ability. Ecosphere, 10, e02783.

Lamie, E., Morton, E.R. & Parzer, H.F. (2025) Puddling in butterflies: current knowledge and new directions. Annals of the Entomological Society of America, 118, 110–118.

Li, H., Zhu, J., Cheng, Y., Zhuo, F., Liu, Y., Huang, J., Taylor, B., Luke, B., Wang, M. & González-Moreno, P. (2023) Daily activity patterns and body temperature of the Oriental migratory locust, Locusta migratoria manilensis (Meyen), in natural habitat. Frontiers in Physiology, 14, p.1110998.

Lorrain-Soligon, L., Robin, F., Bertin, X., Jankovic, M., Rousseau, P., Lelong, V. & Brischoux, F. (2023) Long-term trends of salinity in coastal wetlands: Effects of climate, extreme weather events, and sea water level. Environmental Research, 237, 116937.

Pontes, G., Latorre-Estivalis, J.M., Gutiérrez, M.L., Cano, A., Berón De Astrada, M., Lorenzo, M.G. & Barrozo, R.B. (2022) Molecular and functional basis of high-salt avoidance in a blood-sucking insect. iScience, 25, 104502.

Ramasamy, R., Thiruchenthooran, V., Jayadas, T.T.P., Eswaramohan, T., Santhirasegaram, S., Sivabalakrishnan, K., Naguleswaran, A., Uzest, M., Cayrol, B., Voisin, S.N., Bulet, P. & Surendran, S.N. (2021) Transcriptomic, proteomic and ultrastructural studies on salinity-tolerant Aedes aegypti in the context of rising sea levels and arboviral disease epidemiology. BMC Genomics, 22, 253.

Riedl, C.A., Oster, S., Busto, M., Mackay, T.F. & Sokolowski, M.B. (2016) Natural variability in Drosophila larval and pupal NaCl tolerance. Journal of Insect Physiology, 88, 15–23.

Santiago-Rosario, L.Y., Harms, K.E. & Craven, D. (2022) Contrasts among cationic phytochemical landscapes in the southern United States. Plant Environment Interactions, 3, 226–241.

Santiago-Rosario, L.Y., Shephard, A.M., Snell-Rood, E., Herrmann, A.D. & Harms, K.E. (2025) Butterfly species vary in sex-specific sodium accumulation from larval diets. Ecological Entomology, 50, 228–234.

Silver, S. & A. Donini. (2021) Physiological responses of freshwater insects to salinity: molecular-, cellular- and organ-level studies. Journal of Experimental Biology, 224, jeb222190.

Snell-Rood, E.C., Espeset, A., Boser, C.J., White, W.A. & Smykalski, R. (2014) Anthropogenic changes in sodium affect neural and muscle development in butterflies. Proceedings of the National Academy of Sciences of the USA, 111, 10221–10226.

Vasquez, E.A., Glenn, E.P., Guntenspergen, G.R., Brown, J.J. & Nelson, S.G. (2006) Salt tolerance and osmotic adjustment of Spartina alterniflora (Poaceae) and the invasive M haplotype of Phragmites australis (Poaceae) along a salinity gradient. American Journal of Botany, 93, 1784–1790.

Vincent, S.E. (2006) Sex-based divergence in head shape and diet in the Eastern lubber grasshopper (Romalea microptera). Zoology, 109, 331–338.

Yang, L., Ravikanthachari, N., Marino-Perez, R., Deshmukh, R., Wu, M., Rosenstein, A., Kunte, K., Song, H. & Andolfatto, P. (2019) Predictability in the evolution of Orthopteran cardenolide insensitivity. Philosophical Transactions of the Royal Society of London B Biological Sciences, 374, 20180246.

Yee, D. A., Dean, C., Webb, C., Henke, J.A., Perezchica-Harvey, G., White, G.S., Faraji, A., Macaluso, J.D. & Christofferson, R. (2021) No evidence that salt water ingestion kills adult mosquitoes (Diptera: Culicidae). Journal of Medical Entomology, 58, 767–772.

Xiao, K., Shen, K.Zhong, J.-F. & Li, G.-Q. (2010) Effects of dietary sodium on performance, flight and compensation strategies in the cotton bollworm, Helicoverpa armigera (Hübner) (Lepidoptera: Noctuidae). Frontiers in Zoology, 7, 11.

